# Survivin Regulates Intracellular Stiffness and Extracellular Matrix Production in Vascular Smooth Muscle Cells

**DOI:** 10.1101/2022.10.24.513582

**Authors:** Amanda Krajnik, Erik Nimmer, Andra Sullivan, Joseph A. Brazzo, Yuna Heo, Alanna Krug, John Kolega, Su-Jin Heo, Kwonmoo Lee, Brian R. Weil, Deok-Ho Kim, Yongho Bae

## Abstract

Vascular dysfunction is a common cause of cardiovascular diseases characterized by the narrowing and stiffening of arteries, such as atherosclerosis, restenosis, and hypertension. Arterial narrowing results from the aberrant proliferation of vascular smooth muscle cells (VSMCs) and their increased synthesis and deposition of extracellular matrix (ECM) proteins. These, in turn, are modulated by arterial stiffness, but the mechanism for this is not fully understood. We found that survivin (an inhibitor of apoptosis) is an important regulator of stiffness-mediated ECM synthesis and intracellular stiffness in VSMCs. Whole-transcriptome analysis and cell culture experiments showed that survivin expression is upregulated in injured femoral arteries in mice and in human VSMCs cultured on stiff fibronectin-coated hydrogels. Suppressed expression of survivin in human VSMCs and mouse embryonic fibroblasts decreased the stiffness-mediated expression of ECM components implicated in arterial stiffness, namely, collagen-I, fibronectin, and lysyl oxidase. By contrast, expression of these proteins was upregulated by the overexpression of survivin in human VSMCs cultured on soft hydrogels. Atomic force microscopy analysis showed that suppressed or enhanced expression of survivin decreases or increases intracellular stiffness, respectively. These findings suggest a novel mechanism by which survivin modulates arterial stiffness.

## INTRODUCTION

Vascular smooth muscle cells (VSMCs) are a major component of arterial wall structure and comprise the mechanically active cell layer in the tunica media. Vascular injury triggers many changes to VSMCs, including increased production of extracellular matrix (ECM) components [1–7], including type 1 collagen (collagen-1), fibronectin, and lysyl oxidase (Lox) [8–12]. These changes result in arterial stiffening and can lead to neointima formation and thickening [13–17], which ultimately promote cardiovascular disease [18].

Stiffened arteries in conditions such as vascular injury, atherosclerosis, and hypertension also exhibit an upregulation of an evolutionarily conserved protein called survivin [19–21], also known as baculoviral inhibitor of apoptosis repeat-containing protein 5 (BIRC5) [22]. The increase is attributable to VSMCs, because VSMCs derived from stiffened arteries of spontaneously hypertensive rats produce more survivin, collagen, and fibronectin than those from control rats [23]. However, the mechanism for these changes is not clear.

ECM stiffness is an important mechanical signal for the modulation of cellular function and molecular signaling in VSMCs [24–29], and a stiffer microenvironment is associated with cardiovascular disease [30–33]. Furthermore, mechanical force is an essential component of ECM interactions [34, 35], and the interactions between cells and their substrate lead to altered cell morphology, contractility, and stiffness [36–39], which are crucial for cell adhesion, migration, proliferation, and survival [36, 40–43]. Wall shear stress is associated with the development of intimal hyperplasia and also triggers increased survivin expression and collagen accumulation in VSMCs [44]. Therefore, we sought to explore how the stiffness of the microenvironment signals changes to ECM production in VSMCs and whether survivin is a regulator of this process.

## RESULTS

### Genome-wide analysis identifies vascular injury-related expression of ECM components and survivin

We performed a functional enrichment analysis on differentially expressed genes (DEGs) from previously published microarray data [45] comparing samples of injured and uninjured mouse femoral arteries. The 25,253 genes in the database were filtered for *q* values of ≤0.15 and fold changes of ≥2.0, revealing a total of 667 DEGs: 332 upregulated and 335 downregulated. These DEGs were further analyzed based on the Gene Ontology (GO) database using g:Profiler (https://biit.cs.ut.ee/gprofiler). VSMCs at the site of vascular injury in mice were enriched for genes that code for ECM organization/remodeling and cell surface signaling, including those for integrin, collagen, and fibronectin (**Fig. 1A**).

**Figure 1:**
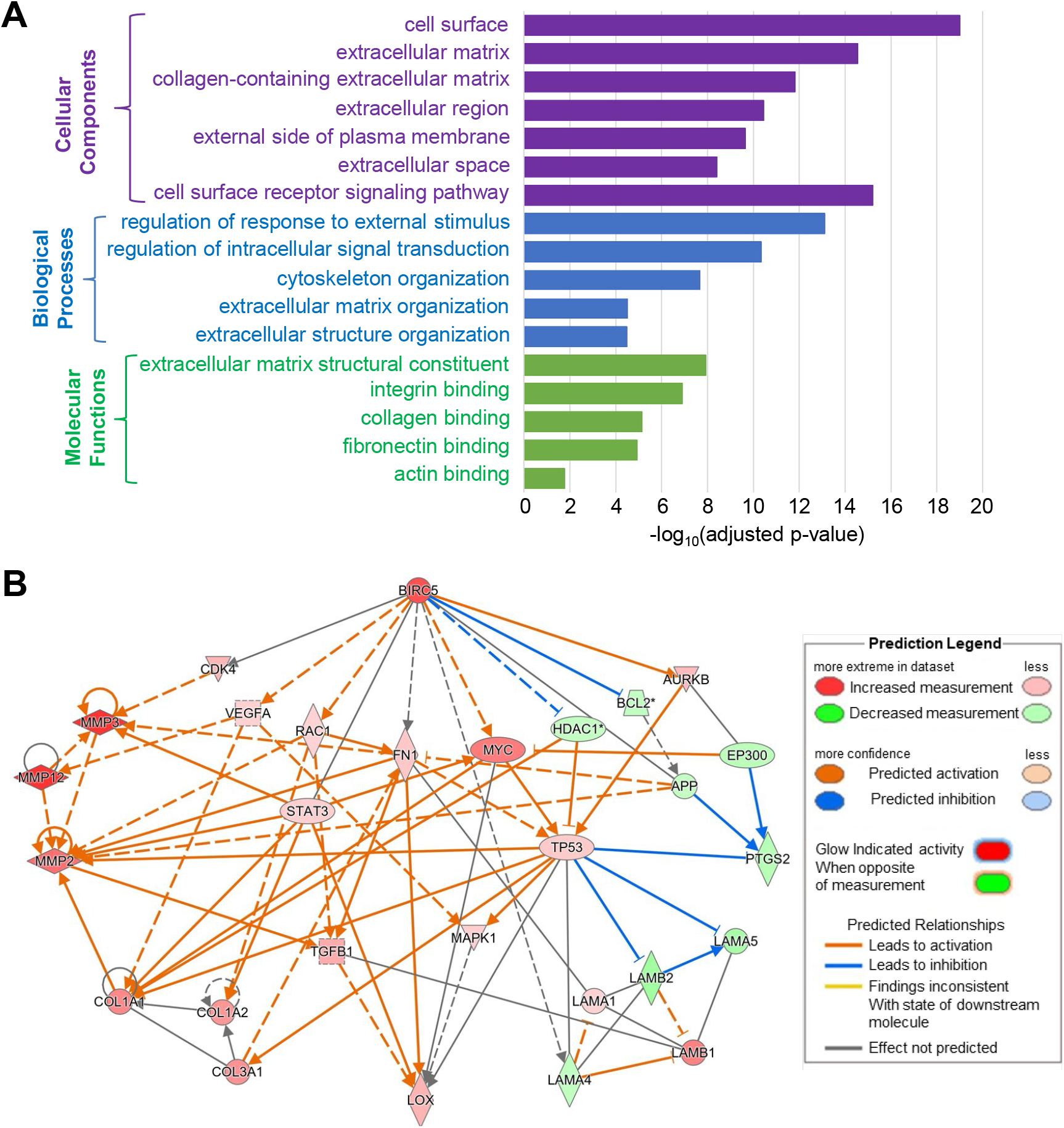
Functional network analysis and predicted network of survivin (*Birc5*) in VSMCs from injured mouse arteries. (**A**) Histogram presenting the significant ECM biological processes (purple), cellular components (blue), and molecular functions (green) enriched among the 332 DEG upregulated in mouse arteries after injury. (**B**) Network diagram of gene interaction pathways between *Birc5* (survivin) and various ECM proteins, including collagens (*Col1a1*, *Col1a2*, and *Col3a1*), fibronectin (*Fn1*), and lysyl oxidase (*Lox*).

To identify the relationship between survivin and these DEGs, we used Ingenuity Pathway Analysis (IPA) to build a potential mechanistic network with predictions based on the DEGs. We use the “my pathway” tool to show connections between *Birc5* (the gene encoding survivin) and all other ECM-related DEGs (**Fig. 1B**). The network illustrates the associations between *Birc5* and genes for ECM proteins such as collagens (*Col1a1*, *Col1a2*, and *Col3a1*), fibronectin (*Fn1*), and lysyl oxidase (*Lox*) in injured arteries. These analyses demonstrate a potential link between survivin and the ECM in vascular injury that can lead to arterial stiffening.

### ECM stiffness modulates survivin expression

To investigate the survivin-ECM link, we examined the expression levels of survivin in VSMCs cultured on soft (2–4 kPa) and stiff (12–25 kPa) fibronectin-coated polyacrylamide hydrogels that approximate the physiological stiffness of healthy and diseased vessels, respectively [45, 46]. For these studies, human VSMCs (hVSMCs) were synchronized in G_0_ by serum starvation and plated on the hydrogels with 10% fetal bovine serum (FBS). After 24 h, the cells were collected for reverse transcription-quantitative PCR (RT-qPCR) and immunoblotting assays. Cells grown on stiff hydrogels had increased expression of survivin at the mRNA (**Fig. 2A**) and protein (**Fig. 2E**) levels compared to that in cells grown on the soft hydrogels (see also **Fig. S1A, B**). hVSMCs plated on stiff hydrogels also had increased expression of three major ECM components, namely collagen-1 (**Fig. 2B, F**), fibronectin (**Fig. 2C, G**), and Lox (**Fig. 2D, H**). Similar results were obtained from mouse embryonic fibroblasts (MEFs) as increased expression levels of survivin (**Fig. S1C, D and S2A**) and ECM components (**Fig. S2B, C**) were observed in MEFs cultured on stiff hydrogels vs. MEFs cultured on soft hydrogels.

**Figure 2:**
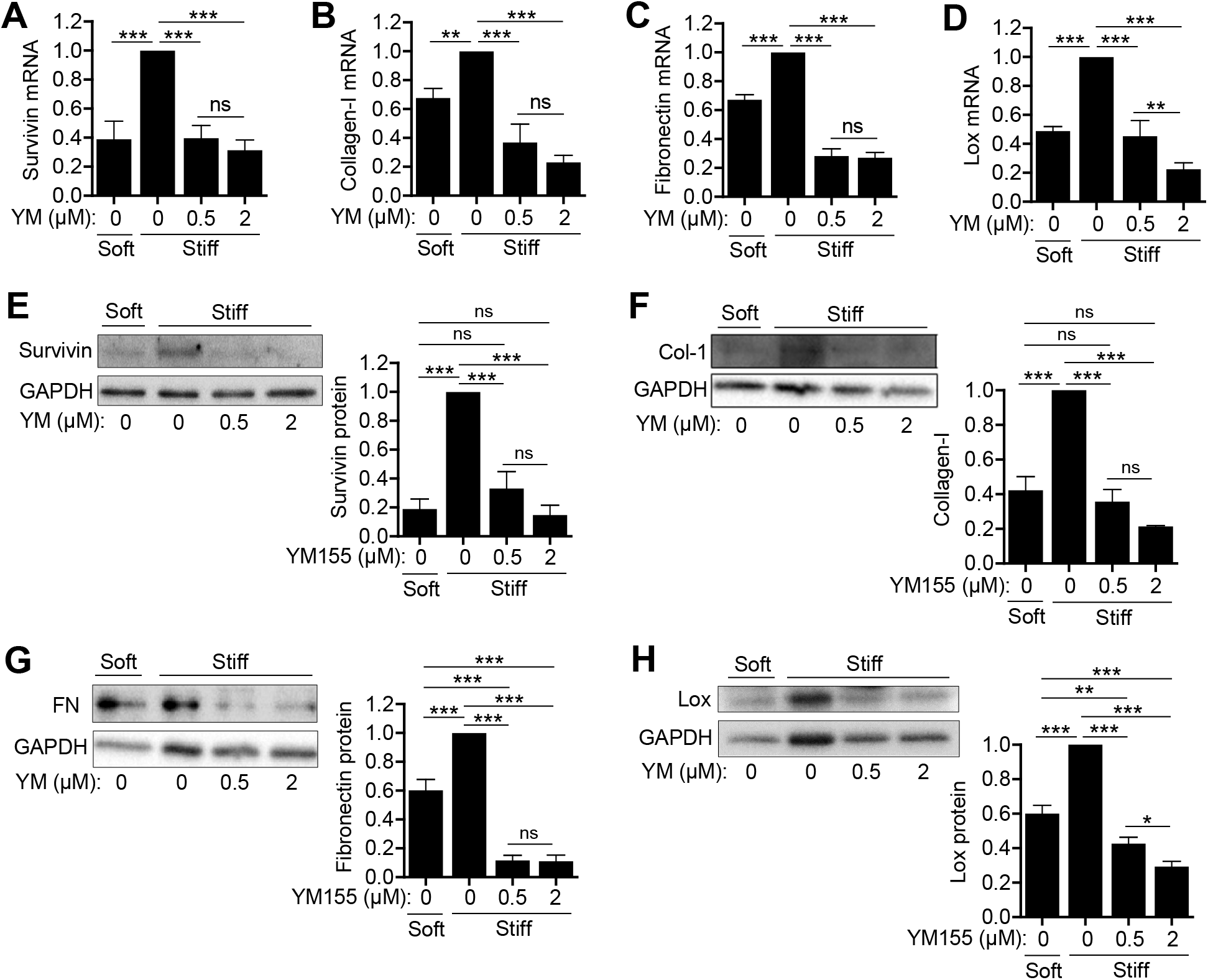
ECM synthesis in hVSMCs is reduced when survivin expression is suppressed. hVSMCs were plated on soft or stiff hydrogels with 10% FBS ± YM155 at the indicated concentrations for 24 h. Levels of mRNA (**A–D**) and protein (**E−H**) were analyzed by RT-qPCR and immunoblotting assays, respectively. The graphs show expression of survivin (**A, E**), collagen-I (**B, F**), fibronectin (**C, G**), and Lox (**D, H**). Expression was normalized to that in hVSMCs treated with DMSO (vehicle control) on stiff hydrogels. *n* = 3−8 (A−D), *n* = 4 (E, H), *n* = 3 (F, G). Error bars show SEMs. *\*p* < 0.05; *\*\*p* < 0.01; *\*\*\*p* < 0.001; ns, not significant by ANOVA followed by Newman–Keuls post hoc test for multiple comparisons.

To identify the role of survivin in the changes observed in hVSMCs grown on stiff substrates, we treated cells with sepantronium bromide (YM155), which suppresses survivin expression, and measured the expression of several ECM components. YM155 at a concentration of 0.5 μM was sufficient to reduce survivin mRNA and protein levels to those seen in cells grown on the soft substrate (**Fig. 2A, E**). Furthermore, YM155 blocked the induction of collagen-I (**Fig. 2B, F**), fibronectin (**Fig. 2C, G**), and Lox (**Fig. 2D, H**), reducing expression to levels below that observed in cells grown on soft substrates. YM155 also reduced the transcript levels of these ECM proteins in MEFs (**Fig. S2**).

We next assessed the effect of survivin overexpression. hVSMCs were infected with adenovirus encoding wild-type survivin (wt-Svv) (multiplicity of infection [MOI] of 25 or 50) or green fluorescent protein (GFP) as the control (MOI, 50) and plated on soft and stiff hydrogels for 24 h. In hVSMCs cultured on the soft substrate, infection with wt-Svv at an MOI of 25 was sufficient to induce survivin levels to those observed in cells cultured on the stiff substrate, with even greater induction seen at an MOI of 50 (**Fig. 3A**). Similarly, wt-Svv induced production of collagen, fibronectin (**Fig. 3B**), and Lox (**Fig. 3C**) to levels observed in cells grown on stiff substrates, with greater induction at the higher MOI. These results show that survivin is sufficient to mimic the ECM production induced when cells are cultured on a stiff substrate. Taken together, the findings suggest that survivin responds to stiffness by coordinating the production of ECM proteins.

**Figure 3:**
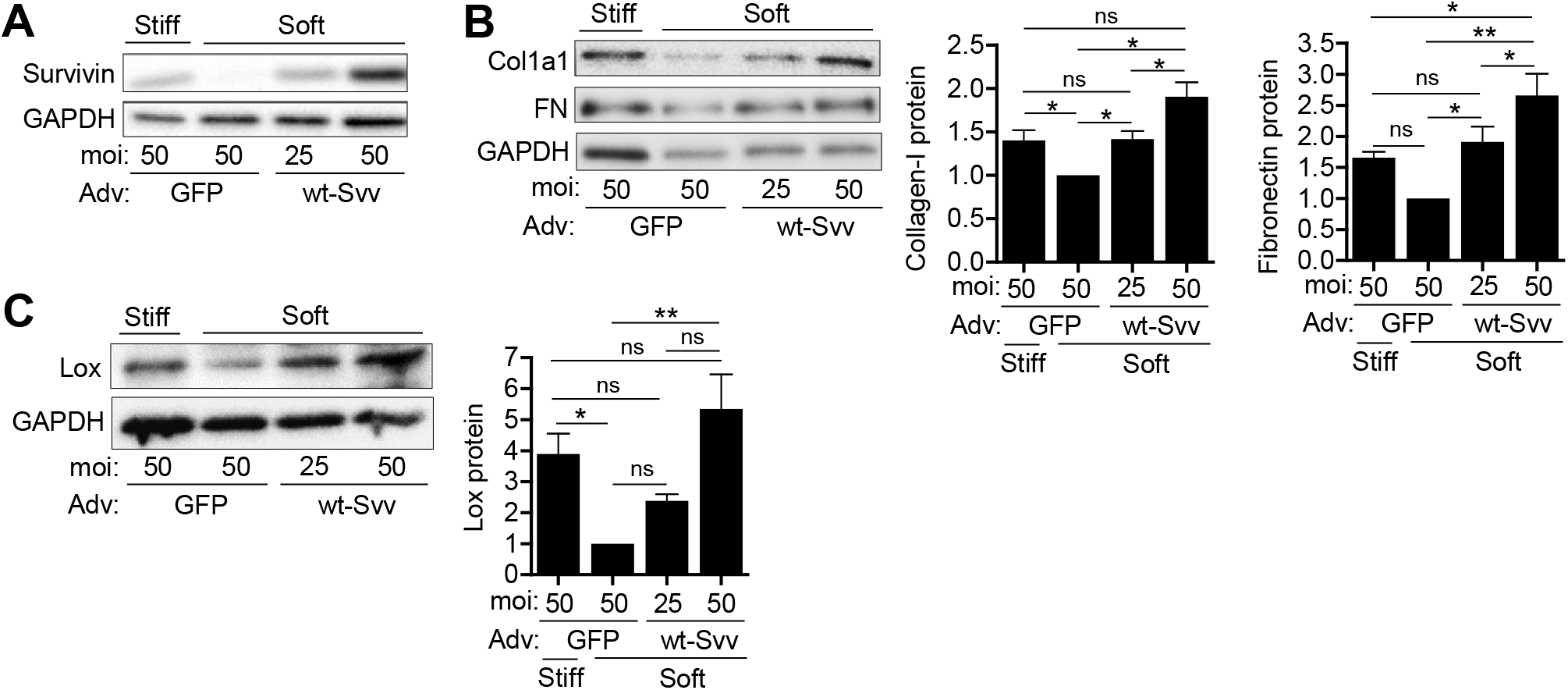
Survivin overexpression mimics stiffness-mediated ECM protein production in hVSMCs. hVSMCs infected with adenoviruses encoding wild-type survivin (wt-Svv) or the GFP control were plated on soft or stiff hydrogels with 10% FBS for 24 h. Total cell lysates were analyzed by immunoblotting for survivin (**A**) collagen, fibronectin (**B**), and Lox (**C**). Levels were normalized to those in GFP-expressing hVSMCs on soft hydrogels. *n* = 5 (A), *n* = 3–4 (B), *n* = 4 (C). Error bars show SEMs. *\*p* < 0.05; *\*\*p* < 0.01; ns, not significant by ANOVA followed by Newman–Keuls post hoc test for multiple comparisons.

### ECM stiffness triggers intracellular stiffness via survivin

In addition to increased ECM protein production, a stiff substrate affects the mechanical properties of VSMCs, i.e., intracellular stiffness and traction force [36, 47–50]. Thus, we explored the role of survivin in ECM stiffness-mediated changes to intracellular stiffness. We performed atomic force microscopy to measure the intracellular stiffness of hVSMCs (**Fig. 4A**) and confirmed that the intracellular stiffness is increased when cells are grown on a stiff substrate, i.e., the stiff fibronectin-coated hydrogel. This increase in intracellular stiffness was blocked when cells were treated with YM155 to reduce survivin expression (**Fig. 4B**). Similar results were obtained with MEFs treated with YM155 (**Fig. S3**). Lastly, overexpression of survivin was sufficient to increase the stiffness of VSMCs plated on a soft hydrogel (**Fig. 4C**). These observations indicate that survivin can regulate intracellular stiffness.

**Figure 4:**
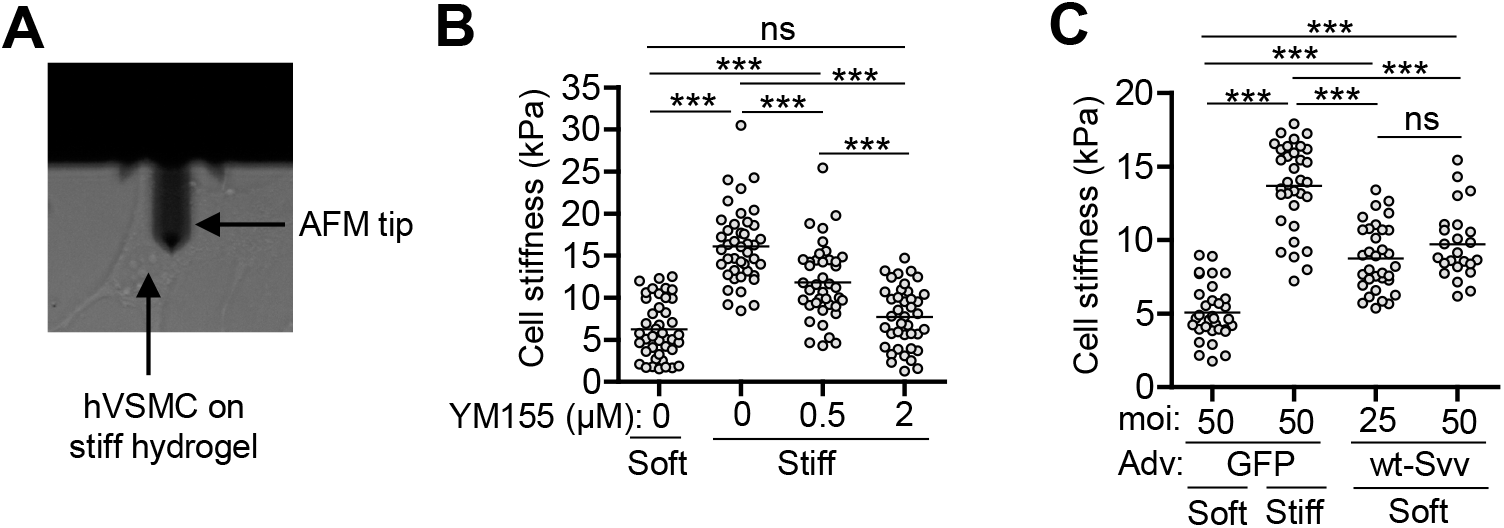
Survivin regulates VSMC stiffness. **(A)** Visualization of an AFM tip on VSMC surface. hVSMCs were plated on fibronectin-coated soft or stiff hydrogels with 10% FBS for 24 h. Atomic force microscopy was used to measure cellular stiffness in cells treated with YM155 (**B**) to reduce survivin levels and in cells infected with adenovirus encoding wild-type survivin (wt-Svv) or a GFP control for survivin overexpression (**C**). *n* = 40−42 cells from four experiments (B) and *n* = 24−35 cells from three experiments (C). Each data point represents one cell, and the means are indicated by horizontal lines. \*\*\**p* < 0.001; ns, not significant by ANOVA followed by Newman– Keuls post hoc test for multiple comparisons.

### Survivin regulates stiffness-mediated Cox2 expression

The data presented above indicate that survivin is an important regulator of ECM production and intracellular stiffness in VSMCs. However, it is not yet clear how survivin enacts this regulation. Our previous studies showed that the expression pattern of cyclooxygenase-2 (Cox2), an enzyme involved in prostaglandin biosynthesis, is the inverse of that for ECM proteins in VSMCs on stiff and soft substrates [51, 52]. Therefore, we reasoned that Cox2 signaling may be involved with the regulatory effects of survivin in VSMCs. We used IPA to build a gene interaction network for *Ptgs2*, the gene encoding Cox2 (also known as prostaglandin-endoperoxide synthase 2), and other ECM genes. On the basis of the directionality of the gene interaction arrows and gene expression levels, the network provides evidence that Cox2 contributes to the regulation of ECM proteins (**Fig. 5A**).

**Figure 5:**
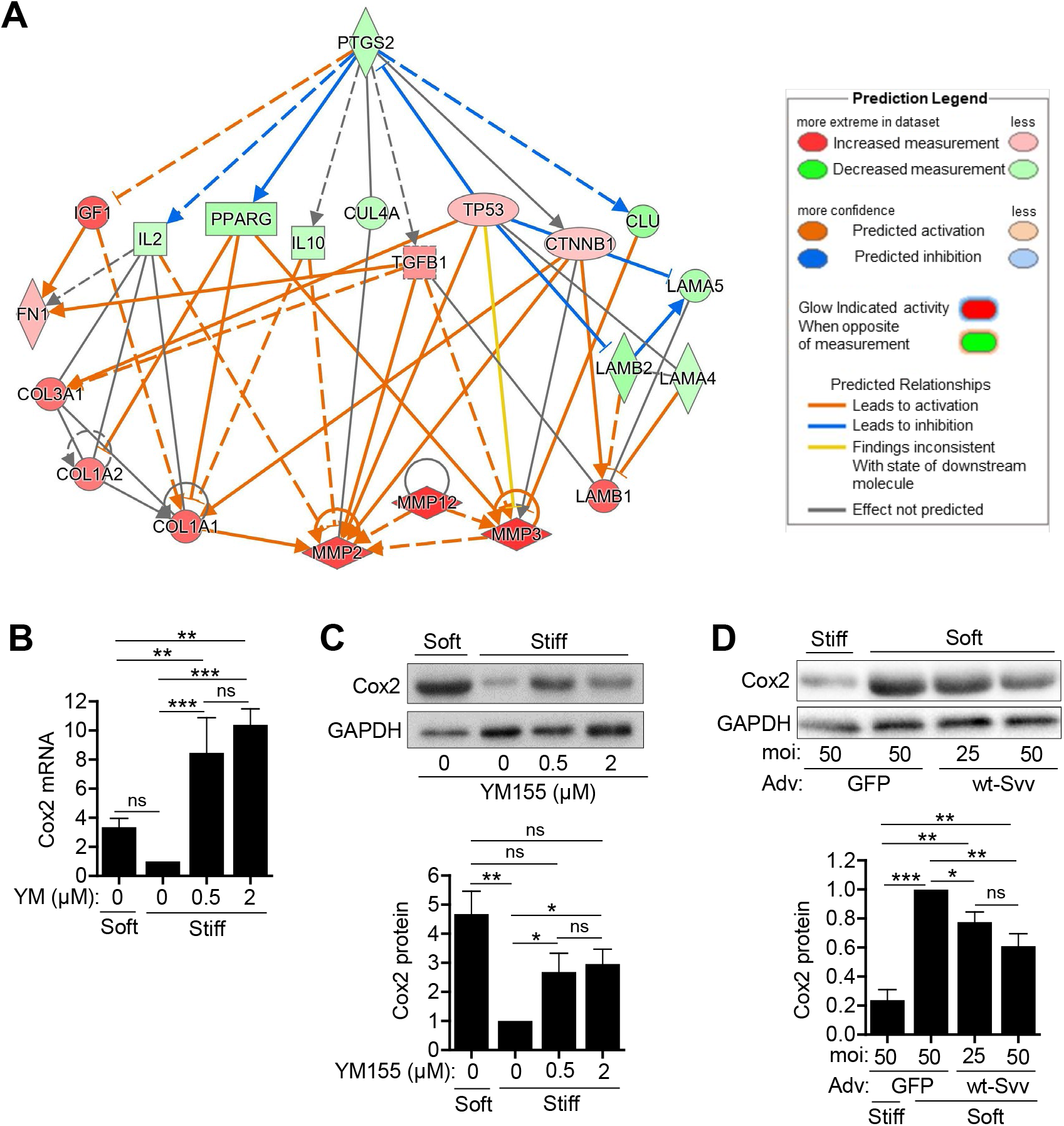
Survivin regulates stiffness-mediated Cox2 expression in hVSMCs. (**A**) Network diagrams of gene interaction pathways between *Ptgs2* (Cox2) and genes for various ECM proteins. hVSMCs were plated on fibronectin-coated soft or stiff hydrogels with 10% FBS ± YM155 at the indicated concentrations for 24 h. Total cell lysates were analyzed by RT-qPCR (**B**) and immunoblotting assays (**C**). hVSMCs infected with adenoviruses encoding GFP or wild-type survivin (wt-Svv) were plated on hydrogels with 10% FBS for 24 h. Cell lysates were analyzed by immunoblotting (**D**). Expression levels were normalized to hVSMCs treated with DMSO (vehicle control) on stiff hydrogels or infected with the virus encoding GFP at an MOI of 50 and plated on soft gels. *n* = 5 (B), *n* = 3 (C, D). GAPDH served as a loading control. Error bars show SEMs. *\*p* < 0.05; *\*\*p* < 0.01; *\*\*\*p* < 0.001; ns, not significant by ANOVA followed by Newman–Keuls post hoc test for multiple comparisons.

We then measured the expression of Cox2 in hVSMCs plated on fibronectin-coated hydrogels. Cox2 was expressed at low levels in cells on stiff substrates, but its expression was markedly upregulated at the mRNA (**Fig. 5B**) and protein (**Fig. 5C**) levels when the cells were treated with YM155 to reduce survivin expression. YM155 treatment of MEFs on stiff substrates restored Cox2 expression to levels observed in cells on soft substrates (**Fig. S4**). Furthermore, we found that survivin overexpression decreased Cox2 levels in hVSMCs plated on soft hydrogels (**Fig. 5D**). Together with the IPA analysis, these results implicate Cox2 in the regulation of ECM proteins and indicate that survivin is a regulator of this function.

## DISCUSSION

Survivin is expressed in response to vascular injury, atherosclerosis, and hypertension in animal models [19, 20, 53] and in proliferating VSMCs in the neointima and media in human atherosclerotic plaques and stenotic vein grafts [20]. However, the role of survivin and the mechanism of its regulation are unknown. We found that survivin expression is sensitive to the stiffness of the ECM and that it induces the production of ECM proteins. Thus, survivin is not only important under conditions associated with arterial stiffening but also possibly exacerbates the stiffening process.

The increased arterial stiffness caused by neointima formation alters the mechanical environment of VSMCs [54, 55]. ECM stiffness and mechanical signals regulate key cellular processes in cardiovascular biology and disease [56]. Blanc-Brude et al. [19] found that survivin is critical for VSMC survival *in vivo* after acute vascular injury, which is associated with arterial stiffening, suggesting survivin may be stiffness sensitive [19]. Our results support this, because survivin was upregulated in cells plated on stiff hydrogels. Our data also suggest that survivin is critical in the transduction of ECM stiffness into intracellular stiffness, which occurs between transmembrane receptors, integrins, and their associated focal adhesion proteins and the actin cytoskeleton [34, 45, 57–60].

The expression of the wild-type survivin was sufficient to induce ECM production on soft substrates. Furthermore, the stiffness-sensitive induction of ECM proteins was blocked when survivin expression was suppressed, indicating that survivin may serve as a signal to promote ECM production. These findings are consistent with other studies showing that downregulation of survivin reduces vessel wall thickness and neointima formation in response to vascular injury [61] and reduces collagen-1 in human Tenon’s capsule fibroblasts [49] and hepatic stellate cells [62]. We posit that survivin expression stimulates ECM synthesis through two distinct stiffness-dependent pathways: one via Lox activation and the other via Cox2 inhibition. Lox is a collagen crosslinker in the ECM, and its expression paralleled that of survivin. Lox deletion or pharmacological inhibition in mice reduces arterial stiffness and collagen accumulation in atherosclerotic plaques [52, 63, 64]. Cox2 also plays a role in cardiovascular biology [65, 66], possibly contributing to the maintenance of healthy vessels by downregulating collagen and fibronectin synthesis [51, 52]. We found that Cox2 expression is inversely related to both ECM stiffness and survivin expression, and understanding this relationship will give more insight into the molecular processes involved in arterial stiffness. Our data indicate that Cox2 expression is inhibited by survivin and associated with the suppression of ECM production. This new finding is significant, because it may provide a strategy to de-stiffen arteries caused by vascular and cardiovascular diseases such as atherosclerosis, coronary artery disease, and hypertension.

## CONCLUSION

Overall, the results of this study showed that survivin expression is stiffness-sensitive and that survivin modulates intracellular stiffness and ECM synthesis in VSMCs. These findings provide new insights into the molecular and mechanical mechanisms that control arterial stiffness and VSMC function as well as potential mechano-therapeutic targets for cardiovascular disease.

## METHODS

### Cell culture

hVSMCs (catalog number [cat. no.] 354-05a, Cell Applications; source: human aorta from a 33-year-old male) were cultured in low-glucose Dulbecco’s modified Eagle’s medium (DMEM) containing 50 μg/ml gentamicin, 1 mM sodium pyruvate, 1× MEM amino acid solution (cat. no. M5550, Sigma-Aldrich) and 10% FBS (cat. no. F2442, Sigma-Aldrich) and used at passages 3–5. Mouse embryonic fibroblasts (MEFs, kind gift of Richard Assoian Laboratory, University of Pennsylvania) were grown in low-glucose DMEM supplemented with 50 μg/ml gentamicin and 10% FBS. Both cell types were maintained in 10% CO_2_ at 37°C. Prior to plating on fibronectin-coated (cat. no. 341631, Calbiochem) polyacrylamide hydrogels, hVSMCs or MEFs near confluence were serum starved for 48 or 24 h, respectively, with DMEM containing 1 mg/ml heat-inactivated, fatty-acid-free bovine serum albumin (BSA; cat. no. 5217, Tocris) to synchronize their cell cycles to G_0_ [45, 46].

### Preparation of fibronectin-coated polyacrylamide hydrogels

The protocol for generating stiffness-tunable polyacrylamide hydrogels was previously described [45, 67]. Glass coverslips, used as bottom coverslips for hydrogel adhesion, were treated with 0.1 M NaOH solution for 5 min to increase the surface area of the coverslip and enable the subsequently added 3-(trimethoxysilyl)propyl methacrylate (cat. no. 440159, Sigma-Aldrich) to attach, to which the fibronectin-coated polyacrylamide hydrogels would be covalently linked. The soft hydrogels (2–4 kPa, mimics the physiological stiffness of a healthy mouse [52, 58]) and stiff hydrogels (12–25 kPa, mimics the physiological stiffness of a diseased mouse artery [52, 58]) were created using various ratios of 40% acrylamide to 1% bis-acrylamide in a solution containing water, 10% ammonium persulfate (cat. no. A3678, Sigma-Aldrich), TEMED (*N*,*N*,*N*′,*N*′-tetramethylethylenediamine; cat. no. J63734.AC, Thermo Scientific) to polymerize the solution, and Tris-fibronectin solution consisting of amine-reactive *N*-hydroxysuccinimide ester (cat. no. A8060, Sigma-Aldrich) dissolved in dimethyl sulfoxide (DMSO; cat. no. D2650, Sigma-Aldrich) added to 1 M Tris-HCl (pH 8.4) with 0.05% fibronectin. This solution was incubated for 2 h at 37°C prior to addition into the hydrogel solution; 150 μl of the prepared solution was used for 24- × 24-mm coverslips, 450 μl was used for 24- × 40-mm coverslips, and 20 μl was used for 12-mm coverslips. Glass coverslips were placed on top of the polyacrylamide hydrogels to spread the solution uniformly across the bottom coverslips. To prevent the hydrogel from attaching to the top coverslips, the coverslips were siliconized with 20% Surfasil (cat. no. TS42801, Thermo Scientific) in chloroform. After polymerization was complete, the top coverslips were removed, and hydrogels were washed three times in 1× Dulbecco’s phosphate-buffered saline (DPBS) for 15 min. Hydrogels were blocked for 30 min in serum-free DMEM with 1% BSA prior to cell plating. For different experiments, different-size glass coverslips and plating densities were used: immunoblotting, 24- × 40-mm coverslips with 1 × 10^5^ cells for stiff hydrogels and 2 × 10^5^ cells for soft hydrogels; RT-qPCR and atomic force microscopy, 24- × 24-mm coverslips with 6 × 10^4^ cells for stiff hydrogels and 1.6 × 10^5^ cells for soft hydrogels; immunostaining, 12-mm coverslips with 1.5 × 10^4^ cells for stiff hydrogels and 3 × 10^4^ cells for soft hydrogels.

### Cell treatments

#### Pharmacological suppression of survivin expression

hVSMCs were serum starved for 48 h with DMEM containing 1 mg/ml BSA and then plated on fibronectin-coated hydrogels for 24 h in DMEM with 10% FBS containing 0.5 μM or 2.0 μM YM155 (an inhibitor of survivin promoter activity; cat. no. 6491, Tocris) in DMSO. MEFs were serum starved for 24 h prior to plating, and YM115 was used at 1.0 μM or 2.0 μM.

#### Adenovirus infection

To overexpress survivin, hVSMCs were incubated in DMEM with 1 mg/ml BSA for ~7 h, and adenovirus harboring *BIRC5* (cat. no. 1611, Vector Biolabs) or the gene for GFP (experimental control; cat. no. 1060, Vector Biolabs) was added to the medium at an MOI of 25 or 50. Cells were serum starved for an additional 40 h before they were plated on fibronectin-coated hydrogels with DMEM containing 10% FBS for 24 h.

### RNA isolation and RT-qPCR

Cells cultured on hydrogels were washed twice with 1× DPBS, and RNA was extracted with TRIzol as described by Thermo Fisher’s TRIzol RNA extraction protocol [68]. A NanoDrop Lite spectrophotometer (cat. no. ND-LITE-PR, Thermo Scientific) was used to determine RNA purity and concentration. Total RNA was reverse transcribed and analyzed by RT-qPCR as previously described [68]. TaqMan probes (Invitrogen) for hVSMCs were used for survivin (*BIRC5;* Hs04194392_m1), collagen-1A1 (*COL1A1*; Hs00164004_m1), fibronectin-1 (*FN1*; Hs01549976_m1), Lox (*LOX*; Hs00942480_m1), Cox2 (*PTGS2*; Hs00153133_m1), and *GAPDH* (Hs02786624_g1). For MEFs, TaqMan probes were used for *Birc5* (Mm00599749_m1), *Col1a1* (Mm00432359_m1), *Lox* (Mm00495386_m1), *Ptgs2* (Mm00478374_m1), and *Gapdh* (Mm99999915_g1). The comparative cycle threshold method was used to determine the mRNA expression for each target gene using the gene for GAPDH as the reference.

### Protein extraction and immunoblotting

hVSMCs cultured on hydrogels were washed twice with cold 1× DPBS. The glass coverslips supporting the hydrogels were placed face-down on 150 μl of 5× sample buffer (250 mM Tris-HCl [pH 6.8], 10% sodium dodecyl sulfate, 50% glycerol, 0.02% bromophenol blue, and 10 mM 2-mercaptoethanol) and incubated for 2 min at room temperature to obtain total cell lysates [45, 46, 67]. The proteins in the resulting total cell lysates were denatured at 100°C, subjected to 6% to 12% sodium dodecyl sulfate-polyacrylamide gel electrophoresis, and transferred electrophoretically onto polyvinylidene difluoride membranes. The membranes were blocked with 6% milk in 1× Tris-buffered saline with 0.1% Tween 20 (TBST) for 1.5 h before they were incubated overnight at 4°C with antibodies to survivin (cat. no. NB500-201, Novus Biologicals), collagen (cat. no. C2456, Sigma-Aldrich), fibronectin (cat. no. F3648, Sigma-Aldrich), Cox2 (cat. no. 66351-1-Ig, Proteintech), Lox (cat. no. NB100-2530, Novus Biologicals), and GAPDH (10494-1-AP, Proteintech). After overnight incubation at 4°C, membranes were washed with 1× TBST for 15 min and probed with secondary antibody for 1 h at room temperature. Blots were washed with 1× TBST for 15 min before imaging. Clarity Western ECL substrate (cat. no. 1705061, Bio-Rad) or Clarity Max Western ECL substrate (cat. no. 1705062, Bio-Rad) were used for antibody detection.

### Atomic force microscopy

Atomic force microscopy was used to measure the intracellular stiffness of cells as described previously [46].The surfaces of the cells cultured on hydrogels were indented with a silicon nitride cantilever (cat. no. BL-AC40TS-C2; Asylum; spring constant, 0.09 N/m) with a three-sided pyramidal tip (8-nm in radius). The stiffness of each cell was measured in contact mode using an NX12 AFM system (Park Systems) mounted on a Nikon ECLIPSE Ti2 inverted microscope. To analyze the stiffness, the first 400 nm of horizontal tip deflection was fit with the Hertz model for a three-sided pyramid and a 35° face angle. For each experimental condition, three force curves per cell were acquired for a total of 10 cells under each condition. Measurements were taken at three evenly spaced locations on the cell membrane. Atomic force microscopy experiments were independently repeated three times (obtaining a total of 30 force curves for each experimental condition). Using atomic force microscopy analysis software XEI (Park Systems), the force curves were quantified and converted to Young’s modulus (stiffness).

### Functional network analysis

#### Gene expression analysis

Differential gene expression analysis was previously performed on microarray data [45]. Duplicate and blank (no name) gene entries were removed, and genes with insignificant differential expression values were filtered out before further analysis. DEGs were defined as those having a fold-change of ≥2.0 and a *q* value of ≤0.15.

#### Functional enrichment analysis

Functional enrichment analysis was performed using the g:GOSt tool in gProfiler (https://biit.cs.ut.ee/gprofiler/gost). The statistical domain scope of the analysis was only annotated genes, and the significance threshold was set to the g:SCS algorithm for computing multiple-testing corrections for *p* values acquired from GO analysis. Significant GO terms were defined by an adjusted *p* value of ≤ 0.05. Biological GO annotations from each of the three main GO categories (Biological Processes, Cellular Components, and Molecular Functions) were considered for this analysis along with those deemed relevant to ECM or mechanosensitive signaling activity. Process GO terms were presented in histograms on a scale of −log(adjusted *p* value).

#### Network analysis

IPA (Qiagen) was used to perform further bioinformatics analysis on the filtered microarray data. A Core Analysis was run, which returned information on various mechanistic pathways and enriched functions on the basis of the literature compiled in the Ingenuity Knowledge Base. The “Diseases and Functions” tool was used to identify molecules known to be involved in ECM synthesis within the microarray dataset, and the “My Pathway” tool was subsequently used to display known relationships between *Birc5* and other genes within the ECM synthesis function. The z-directional components of the expression analysis were based on the expression log ratio values. Functions with a z-score of >2 were regarded to have significant activation, whereas those with a z-score of <−2 were considered to have significant inhibition. The “Molecule Activity Predictor” tool was used to display gene expression levels by node color and intensity and to generate predicted activation states of molecules and interactions based on results of the Core Analysis.

### Statistical analysis

Statistical significance was determined using Prism (Graph-Pad) software. Data are presented as means and standard errors of the means (SEMs) and were analyzed with Student’s *t* tests or ANOVAs followed by Newman–Keuls post hoc test for multiple comparisons as appropriate. Samples sizes for each group are indicated in the figure legends.

## ACKNOWLEDGEMENTS

We thank Karen Dietz for critical reading and editing of the manuscript. This work was supported by American Heart Association Career Development Award 18CDA34080415 and NIH grant 1R56HL163168-01 to Y.B.

## SUPPLEMENTARY FIGURE LEGENDS

**Supplementary Figure S1:**
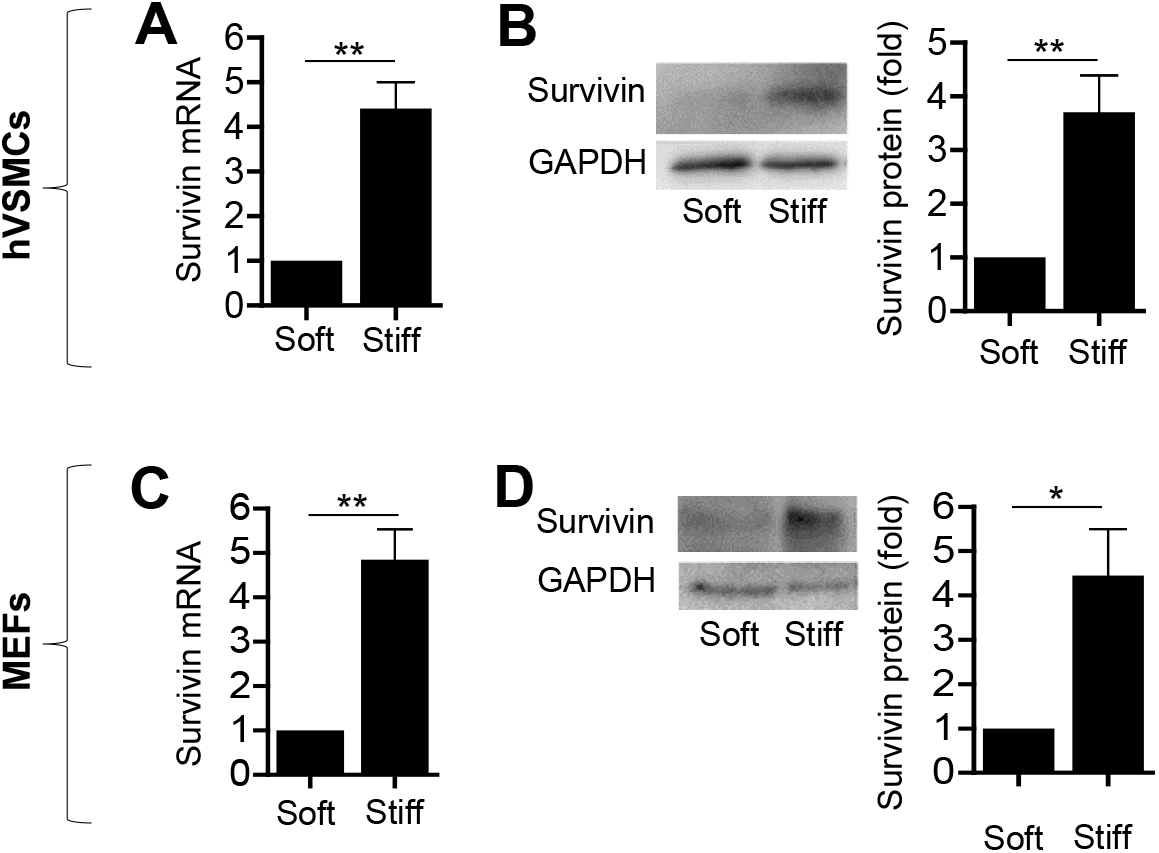
Stiff ECM upregulates the levels of survivin mRNA and protein. Human vascular smooth muscle cells (hVSMCs; **A, B**) and mouse embryonic fibroblasts (MEFs) (**C, D**) were synchronized to G_0_ by serum starvation and plated on fibronectin-coated soft or stiff hydrogels with 10% FBS for 24 h. Total cell lysates were analyzed by RT-qPCR for mRNA (**A, C**) and immunoblotting for protein (**B, D**). Expression levels were normalized to that of GAPDH. *n = 7* (A, C), *n* = 6 (B), *n=4* (D). Error bars show SEMs. *\*p* < 0.05 and *\*\*p* < 0.01 by Student’s *t* test.

**Supplementary Figure S2:**
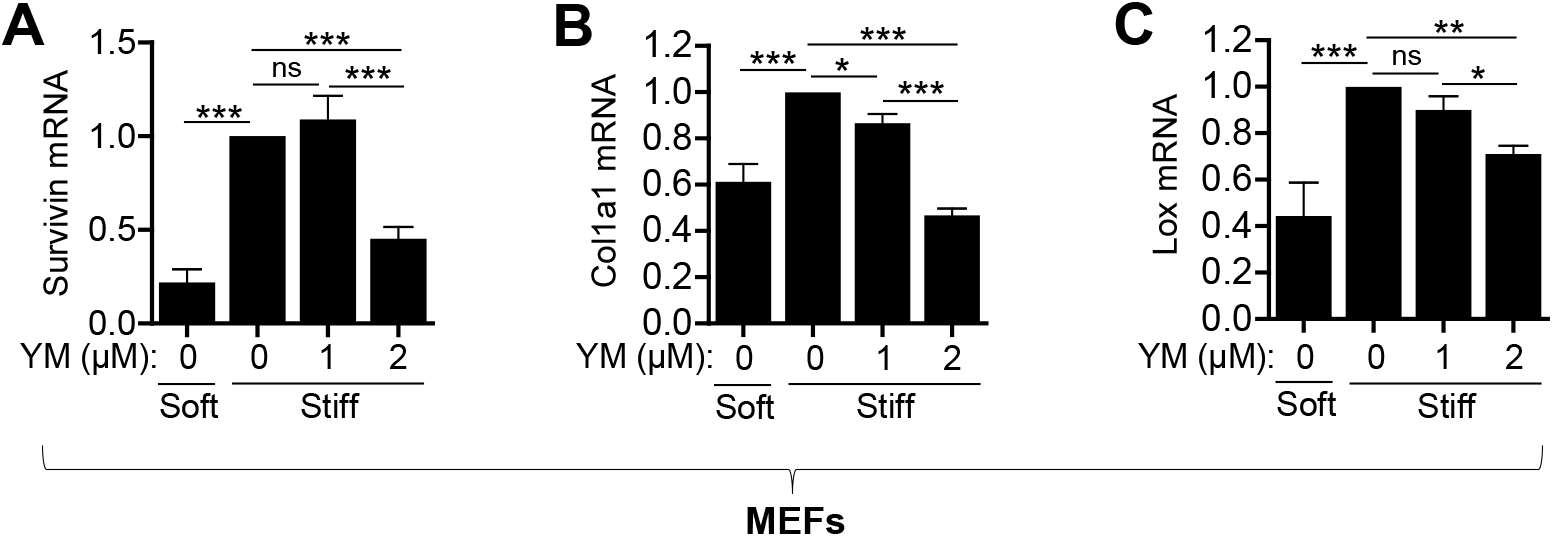
Stiffness-mediated expression of ECM components in mouse embryonic fibroblasts (MEFs) requires survivin. MEFs were plated on soft or stiff hydrogels with 10% FBS ± YM155 at the indicated concentrations for 24 h to suppress survivin expression. RT-qPCR was performed to measure mRNA expression of survivin (**A**), collagen-I (*Col1a1*, **B**), and *Lox* (**C**). Levels were normalized to those in MEFs treated with DMSO (vehicle control) on stiff hydrogels. *n* = 3–5. Error bars show SEMs. \**p* < 0.05; \*\**p* < 0.01; \*\*\**p* < 0.001; ns, not significant by ANOVA followed by Newman-Keuls post hoc test for multiple comparisons.

**Supplementary Figure S3:**
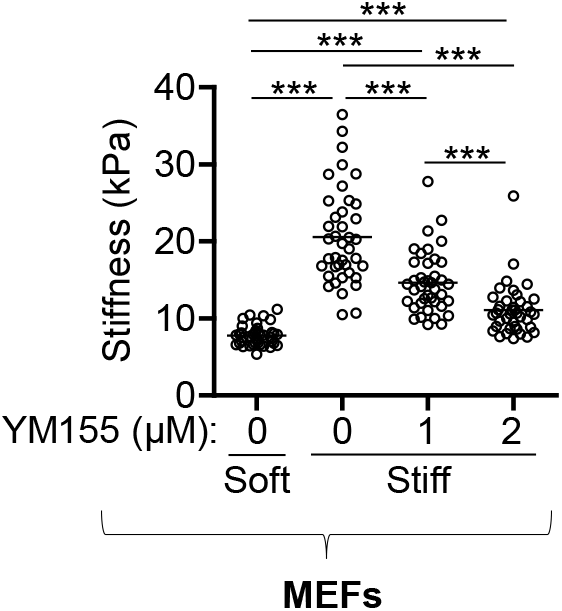
ECM-mediated regulation of intracellular stiffness in MEFs requires survivin. MEFs plated on fibronectin-coated soft or stiff hydrogels with 10% FBS were treated with YM155 at the indicated concentrations to suppress survivin expression for 24 h. Atomic force microscopy was performed to measure intracellular stiffness. *n* = 38−40 cells from three experiments. Each data point represents one cell, and the means are indicated by horizontal lines. \*\*\**p* < 0.001 by ANOVA followed by Newman–Keuls post hoc test for multiple comparisons.

**Supplementary Figure S4:**
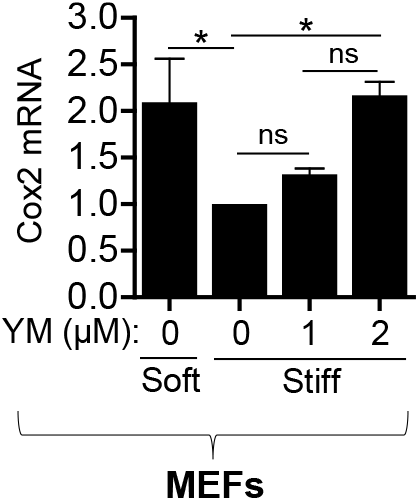
Cox2 expression in MEFs on stiff substrates is upregulated when survivin is suppressed. MEFs were plated on fibronectin-coated soft or stiff hydrogels with 10% FBS ± YM155 at the indicated concentratons for 24 h to reduce survivin expression. mRNA levels were measured via RT-qPCR and normalized to that in MEFs treated with DMSO on stiff hydrogels. *n* = 3. Error bars show SEMs. *\*p* < 0.05; ns, not significant by ANOVA followed by Newman–Keuls post-hoc test for multiple comparisons.

## References

1. Ponticos, M. and B.D. Smith, Extracellular matrix synthesis in vascular disease: hypertension, and atherosclerosis. J Biomed Res, 2014. 28(1): p. 25–39.

2. Feil, S., et al., Transdifferentiation of vascular smooth muscle cells to macrophage-like cells during atherogenesis. Circ Res, 2014. 115(7): p. 662–7.

3. Beamish, J.A., et al., Molecular regulation of contractile smooth muscle cell phenotype: implications for vascular tissue engineering. Tissue Eng Part B Rev, 2010. 16(5): p. 467–91.

4. Owens, G.K., M.S. Kumar, and B.R. Wamhoff, Molecular regulation of vascular smooth muscle cell differentiation in development and disease. Physiol Rev, 2004. 84(3): p. 767–801.

5. Thyberg, J., et al., Regulation of differentiated properties and proliferation of arterial smooth muscle cells. Arteriosclerosis, 1990. 10(6): p. 966–90.

6. Owens, G.K., Regulation of differentiation of vascular smooth muscle cells. Physiol Rev, 1995. 75(3): p. 487–517.

7. Thyberg, J., et al., Phenotypic modulation of smooth muscle cells after arterial injury is associated with changes in the distribution of laminin and fibronectin. J Histochem Cytochem, 1997. 45(6): p. 837–46.

8. von Kleeck, R., et al., Arterial stiffness and cardiac dysfunction in Hutchinson-Gilford Progeria Syndrome corrected by inhibition of lysyl oxidase. Life Sci Alliance, 2021. 4(5).

9. Morawietz, H., LOX-1 and atherosclerosis: proof of concept in LOX-1-knockout mice. Circ Res, 2007. 100(11): p. 1534–6.

10. Martinez-Gonzalez, J., et al., Emerging Roles of Lysyl Oxidases in the Cardiovascular System: New Concepts and Therapeutic Challenges. Biomolecules, 2019. 9(10).

11. Xu, J. and G. P. Shi, Vascular wall extracellular matrix proteins and vascular diseases. Biochim Biophys Acta, 2014. 1842(11): p. 2106–2119.

12. Lacolley, P., et al., Increased carotid wall elastic modulus and fibronectin in aldosterone-salt-treated rats: effects of eplerenone. Circulation, 2002. 106(22): p. 2848–53.

13. Ma, Z., et al., Extracellular matrix dynamics in vascular remodeling. Am J Physiol Cell Physiol, 2020. 319(3): p. C481–C499.

14. Lacolley, P., et al., Vascular Smooth Muscle Cells and Arterial Stiffening: Relevance in Development, Aging, and Disease. Physiol Rev, 2017. 97(4): p. 1555–1617.

15. Zhang, Y., et al., Arterial Stiffness in Hypertension and Function of Large Arteries. Am J Hypertens, 2020. 33(4): p. 291–296.

16. Bonnans, C., J. Chou, and Z. Werb, Remodelling the extracellular matrix in development and disease. Nat Rev Mol Cell Biol, 2014. 15(12): p. 786–801.

17. Sonbol, H.S., Extracellular Matrix Remodeling in Human Disease. J Microsc Ultrastruct, 2018. 6(3): p. 123–128.

18. Pizzolato, R. and J.M. Romero, Neurosonology and noninvasive imaging of the carotid arteries. Handb Clin Neurol, 2016. 135: p. 165–191.

19. Blanc-Brude, O.P., et al., Inhibitor of apoptosis protein survivin regulates vascular injury. Nat Med, 2002. 8(9): p. 987–94.

20. Simosa, H.F., et al., Survivin expression is up-regulated in vascular injury and identifies a distinct cellular phenotype. J Vasc Surg, 2005. 41(4): p. 682–90.

21. Conte, M.S. and D.C. Altieri, Survivin regulation of vascular injury. Trends Cardiovasc Med, 2006. 16(4): p. 114–7.

22. de Almagro, M.C. and D. Vucic, The inhibitor of apoptosis (IAP) proteins are critical regulators of signaling pathways and targets for anti-cancer therapy. Exp Oncol, 2012. 34(3): p. 200–11.

23. Arribas, S.M., et al., Enhanced survival of vascular smooth muscle cells accounts for heightened elastin deposition in arteries of neonatal spontaneously hypertensive rats. Exp Physiol, 2010. 95(4): p. 550–60.

24. Sazonova, O.V., et al., Extracellular matrix presentation modulates vascular smooth muscle cell mechanotransduction. Matrix Biol, 2015. 41: p. 36–43.

25. Brown, X.Q., et al., Effect of substrate stiffness and PDGF on the behavior of vascular smooth muscle cells: implications for atherosclerosis. J Cell Physiol, 2010. 225(1): p. 115–22.

26. Byfield, F.J., et al., Absence of filamin A prevents cells from responding to stiffness gradients on gels coated with collagen but not fibronectin. Biophys J, 2009. 96(12): p. 5095–102.

27. Calve, S. and H.G. Simon, Biochemical and mechanical environment cooperatively regulate skeletal muscle regeneration. FASEB J, 2012. 26(6): p. 2538–45.

28. Sazonova, O.V., et al., Cell-cell interactions mediate the response of vascular smooth muscle cells to substrate stiffness. Biophys J, 2011. 101(3): p. 622–30.

29. Liu, S. and Z. Lin, Vascular Smooth Muscle Cells Mechanosensitive Regulators and Vascular Remodeling. J Vasc Res, 2022. 59(2): p. 90–113.

30. Handorf, A.M., et al., Tissue stiffness dictates development, homeostasis, and disease progression. Organogenesis, 2015. 11(1): p. 1–15.

31. Park, S., et al., The Effects of Stiffness, Fluid Viscosity, and Geometry of Microenvironment in Homeostasis, Aging, and Diseases: A Brief Review. J Biomech Eng, 2020. 142(10).

32. Mitchell, G.F., et al., Arterial stiffness and cardiovascular events: the Framingham Heart Study. Circulation, 2010. 121(4): p. 505–11.

33. Zieman, S.J., V. Melenovsky, and D.A. Kass, Mechanisms, pathophysiology, and therapy of arterial stiffness. Arterioscler Thromb Vasc Biol, 2005. 25(5): p. 932–43.

34. Solon, J., et al., Fibroblast adaptation and stiffness matching to soft elastic substrates. Biophys J, 2007. 93(12): p. 4453–61.

35. Galbraith, C.G., K.M. Yamada, and M.P. Sheetz, The relationship between force and focal complex development. J Cell Biol, 2002. 159(4): p. 695–705.

36. Yeung, T., et al., Effects of substrate stiffness on cell morphology, cytoskeletal structure, and adhesion. Cell Motil Cytoskeleton, 2005. 60(1): p. 24–34.

37. Stroka, K.M. and H. Aranda-Espinoza, Effects of Morphology vs. Cell-Cell Interactions on Endothelial Cell Stiffness. Cell Mol Bioeng, 2011. 4(1): p. 9–27.

38. Doss, B.L., et al., Cell response to substrate rigidity is regulated by active and passive cytoskeletal stress. Proc Natl Acad Sci U S A, 2020. 117(23): p. 12817–12825.

39. Chiang, M.Y., et al., Relationships among cell morphology, intrinsic cell stiffness and cell-substrate interactions. Biomaterials, 2013. 34(38): p. 9754–62.

40. Pelham, R.J., Jr. and Y.L. Wang, Cell locomotion and focal adhesions are regulated by the mechanical properties of the substrate. Biol Bull, 1998. 194(3): p. 348–9; discussion 349–50.

41. Wang, H.B., M. Dembo, and Y.L. Wang, Substrate flexibility regulates growth and apoptosis of normal but not transformed cells. Am J Physiol Cell Physiol, 2000. 279(5): p. C1345–50.

42. Miller, A.E., P. Hu, and T.H. Barker, Feeling Things Out: Bidirectional Signaling of the Cell-ECM Interface, Implications in the Mechanobiology of Cell Spreading, Migration, Proliferation, and Differentiation. Adv Healthc Mater, 2020. 9(8): p. e1901445.

43. Jensen, C. and Y. Teng, Is It Time to Start Transitioning From 2D to 3D Cell Culture? Front Mol Biosci, 2020. 7: p. 33.

44. Zhang, H., et al., Wall shear stress promotes intimal hyperplasia through the paracrine H2O2-mediated NOX-AKT-SVV axis. Life Sci, 2018. 207: p. 61–71.

45. Bae, Y.H., et al., A FAK-Cas-Rac-lamellipodin signaling module transduces extracellular matrix stiffness into mechanosensitive cell cycling. Sci Signal, 2014. 7(330): p. ra57.

46. Brazzo, J.A., et al., Mechanosensitive expression of lamellipodin promotes intracellular stiffness, cyclin expression and cell proliferation. J Cell Sci, 2021. 134(12).

47. Bershadsky, A.D., N.Q. Balaban, and B. Geiger, Adhesion-dependent cell mechanosensitivity. Annu Rev Cell Dev Biol, 2003. 19: p. 677–95.

48. Peyton, S.R., et al., The emergence of ECM mechanics and cytoskeletal tension as important regulators of cell function. Cell Biochem Biophys, 2007. 47(2): p. 300–20.

49. Schwartz, M.A., Integrins and extracellular matrix in mechanotransduction. Cold Spring Harb Perspect Biol, 2010. 2(12): p. a005066.

50. Janmey, P.A. and C.A. McCulloch, Cell mechanics: integrating cell responses to mechanical stimuli. Annu Rev Biomed Eng, 2007. 9: p. 1–34.

51. Hsu, B.Y., et al., Apolipoprotein E3 Inhibits Rho to Regulate the Mechanosensitive Expression of Cox2. PLoS One, 2015. 10(6): p. e0128974.

52. Kothapalli, D., et al., Cardiovascular protection by ApoE and ApoE-HDL linked to suppression of ECM gene expression and arterial stiffening. Cell Rep, 2012. 2(5): p. 1259–71.

53. Kobayashi, K., et al., Expression of a murine homologue of the inhibitor of apoptosis protein is related to cell proliferation. Proc Natl Acad Sci U S A, 1999. 96(4): p. 1457–62.

54. Yurdagul, A., Jr., et al., The arterial microenvironment: the where and why of atherosclerosis. Biochem J, 2016. 473(10): p. 1281–95.

55. Wang, J., et al., Endovascular stent-induced alterations in host artery mechanical environments and their roles in stent restenosis and late thrombosis. Regen Biomater, 2018. 5(3): p. 177–187.

56. Wang, G.J., et al., Regulation of vein graft hyperplasia by survivin, an inhibitor of apoptosis protein. Arterioscler Thromb Vasc Biol, 2005. 25(10): p. 2081–7.

57. Xie, J., et al., Energy expenditure during cell spreading influences the cellular response to matrix stiffness. Biomaterials, 2021. 267: p. 120494.

58. Klein, E.A., et al., Cell-cycle control by physiological matrix elasticity and in vivo tissue stiffening. Curr Biol, 2009. 19(18): p. 1511–8.

59. Baker, E.L., R.T. Bonnecaze, and M.H. Zaman, Extracellular matrix stiffness and architecture govern intracellular rheology in cancer. Biophys J, 2009. 97(4): p. 1013–21.

60. Assoian, R.K. and E.A. Klein, Growth control by intracellular tension and extracellular stiffness. Trends Cell Biol, 2008. 18(7): p. 347–52.

61. Wang, X., et al., The Role of Survivin and Transcription Factor FOXP1 in Scarring After Glaucoma Surgery. Transl Vis Sci Technol, 2022. 11(2): p. 19.

62. Sharma, S., et al., Survivin expression is essential for early activation of hepatic stellate cells and fibrosis progression in chronic liver injury. Life Sci, 2021. 287: p. 120119.

63. Tian, K., et al., Targeting LOX-1 in atherosclerosis and vasculopathy: current knowledge and future perspectives. Ann N Y Acad Sci, 2019. 1443(1): p. 34–53.

64. Kattoor, A.J., A. Goel, and J.L. Mehta, LOX-1: Regulation, Signaling and Its Role in Atherosclerosis. Antioxidants (Basel), 2019. 8(7).

65. Ali, K., et al., Structure-function properties of the apoE-dependent COX-2 pathway in vascular smooth muscle cells. Atherosclerosis, 2008. 196(1): p. 201–209.

66. Grosser, T., S. Fries, and G.A. FitzGerald, Biological basis for the cardiovascular consequences of COX-2 inhibition: therapeutic challenges and opportunities. J Clin Invest, 2006. 116(1): p. 4–15.

67. Klein, E.A., et al., Cell adhesion, cellular tension, and cell cycle control. Methods Enzymol, 2007. 426: p. 155–75.

68. Oudit, G.Y., et al., Loss of PTEN attenuates the development of pathological hypertrophy and heart failure in response to biomechanical stress. Cardiovasc Res, 2008. 78(3): p. 505–14.

